# Robustness of bidirectional microtubule network self-organization

**DOI:** 10.1101/825786

**Authors:** Aleksandra Z. Płochocka, Alexander M. Davie, Natalia. A. Bulgakova, Lyubov Chumakova

**Affiliations:** Center for Computational Biology and Center for Computational Mathematics, Flatiron Institute, New York, NY, USA, 10010; Maxwell Institute for Mathematical Sciences, School of Mathematics, The University of Edinburgh, Edinburgh, UK, EH9 3FD; Department of Biomedical Science, The University of Sheffield, Sheffield, UK, S10 2TN

**Keywords:** cell biology, microtubule, self-organization, robustness

## Abstract

Robustness of biological systems is crucial for their survival, however, for many systems its origin is an open question. Here we analyze one sub-cellular level system, the microtubule cytoskeleton. Microtubules self-organize into a network, along which cellular components are delivered to their biologically relevant locations. While individual microtubule are highly dynamic with their dynamics depends on the organism environment and genetics, network sensitivity to this dynamics would result in pathologies. Combining mathematical modelling with genetic manipulations in *Drosophila*, we show that the microtubule self-organization indeed does not depend on dynamics of individual microtubules, and thus is robust on the tissue level. We demonstrate the origin of this robustness via a mathematical model, suggesting this being a generic mechanism.

## I. INTRODUCTION

The correct positioning of intracellular components such as proteins and organelles is critical for correct cellular function [1, 2]. These components are transported to their biologically relevant locations by motor proteins moving along the cytoskeleton [3–5]. Therefore, the direction of cytoskeleton filaments guides the direction and efficiency of intracellular transport. One common type of cytoskeleton used for transport is microtubules [3]. These are highly dynamic unstable polymers which switch between periods of growth and shrinkage. During the growth phase, GTP-tubulin dimers are added to the microtubule plus-end forming a GTP-cap. GTP-tubulin stochastically hydrolizes into GDP-tubulin, and the loss of the GTP-cap results in a catastrophe: a switch to depolymerization [6]. Microtubule dynamical properties are influenced by a wide range of both internal and external factors. For example, the dynamics and number of individual microtubules in a cell depend on the expression of particular plus- and minus-end binding proteins [7]; interaction between microtubules is affected by the presence of cross-linking and motor proteins [8]; and the stability of the microtubule network is affected by external factors, e.g. temperature, which changes microtubule rigidity [9].

Animal cells take multiple shapes and forms depending on their function, ranging from neuronal cells with meter-long projections, to epithelial cells, to migrating ameboidal leukocytes. Therefore, it is not surprising that microtubule systems similarly acquire a multitude of organizations. In undifferentiated cells, microtubules form a radial array with minus-ends at a microtubule organizing centre at a centrosome. Upon cell differentiation, microtubules are reorganized into non-centrosomal arrays of varying geometry, ranging from unidirectional bundles (e.g. axons), to bidirectional microtubule systems (e.g. subapical microtubules or microtubules in dendrites) [10]. The microtubule self-organization in unidirectional or radial microtubule systems has been extensively studied, for example, [8, 11–13]. Directionality and alignment of these networks depend on several interconnected factors. These include the localization of microtubule minus-ends, which could be all concentrated at a single location (microtubule organizing centre), distributed uniformly at the cell surface, or targeted to specific locations, for example to the sites of cell-cell contacts [10]. Other factors are the geometrical constrains of a cell, for example microtubules can grow only in specific directions in a long and thin axon or dendrite, and the presence of cross-linking and motor proteins. Bidirectional microtubule networks have an additional level of complexity, as several other factors independently contribute to their organization: the dynamics of individual microtubules, their interaction, and the often non-trivial distribution of minus-ends.

In this paper we explore the self-organization of bidirectional microtubule networks, which we define as the degree of alignment of individual microtubule filaments with each other. Such networks are particularly common in differentiated epithelial cells, which form one of the four fundamental tissue types found in all animals [10, 14–16]. These networks are constrained in space forming a quasi-2d system, which enable modelling it using a 2d mathematical model [17]. In these cells microtubules form a dense mesh just under the apical surface, and are seeded from sites of cell-cell adhesion at the cell periphery [18]. Note, here we only consider mixed-orientation microtubules within the apical plane, and not unidirectional apical-basal microtubules. A systematic analysis of microtubule self-organization dependence on each parameter is prohibitive *in vivo*, since the number of combinations of individual dependencies is extremely large and goes beyond the microtubule network itself. In addition, altering the microtubule network has profound consequences on processes relying on them, and thus, on cellular functions. We therefore analyze this system via mathematical modelling, and validate the model predictions *in vivo* when possible.

Various modelling approaches have been used for describing microtubule self-organization. However, many of them are specific to a particular tissue. In plants, it was shown that microtubule zipping strongly affects their self-organization [19]; and that tension can have a non-negligible effect on stabilizing microtubules [20]. In larger cells such as *Drosophila* oocytes, microtubule nucleation at the cortex was shown to be important [21]. Models that include the hydrodynamic effect of the cytoplasm and molecular motors’ effect on microtubule self-organization are summarized in [22, 23]. Our published stochastic model successfully recapitulates the organization of microtubule networks in various epithelial cells [17]. It is a minimal *in silico* 2d-model, where the microtubules are seeded on the cell periphery, grow stochastically to capture the dynamic instability (as in [24]), and follow geometric interaction rules.

Here, we use this stochastic model for simulations exploring the large parameter space of microtubule dynamics parameterizations, discovering that microtubule self-organization is robust. We confirm the robustness *in vivo* using genetic manipulations of the epithelial cells in the model organism *Drosophila*. Finally, we build a minimal probabilistic model that reveals that the reason for robustness is the separation of time-scales of microtubule dynamics. This model shows that the details of microtubule dynamic instability are irrelevant for microtubule self-organization within their biologically relevant ranges, and that the only biological quality beyond cell shape that affects microtubule alignment is the minus-end distribution. Our minimal model accurately predicts the experimental results given the experimental data. Therefore, we demonstrate extreme robustness of bidirectional quasi-2d microtubule self-organization, which can be explained by simple mathematical rules. This suggests a general applicability of our findings to bidirectional microtubule networks, and provides a foundation for future studies.

## II. RESULTS

### 1. A geometric model accurately predicts in vivo microtubule alignment

Cells of the *Drosophila* epidermis elongate during stages 12-15 of embryonic development, changing their eccentricity from 0.7 to 0.98 (Fig. 1A and [17]). As cells elongate, initially randomly oriented microtubules become gradually aligned [17]. The simplest thought experiment to visualize how cell elongation translates into microtubule alignment is the following. Imagine a “hairy” unit-circle on the (*x, y*) plane, where “hairs” are microtubules. Turn it inside out (Fig. 1B). The micro-tubules are randomly pointing inside the ball, representing absence of microtubule alignment in non-elongated cells; at each microtubule minus-end on the cell boundary, the mean microtubule direction is normal to the cell boundary. Stretching this cell uniformly in the *y*-direction by a factor *b* results in an ellipse with eccentricity 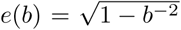. The minus-end position will move proportionally to deformation, whereas the filament direction will point towards the cell center and its length will remain unaltered. Mathematically, the distribution *ρ*(*θ*) of microtubule angles *θ* ∈ [0, 180] changes from the uniform *ρ*(*θ*) = 1*/*180 to the angle distribution we call *hairyball* distribution, *ρ*_*HB*_(*ϕ*), which is the inverse Jacobian of the stretching mapping *F*: *θ → ϕ*, where *θ, ϕ* ∈ [0, 180], given by tan(*ϕ*(*θ*)) = *b* tan(*θ*)

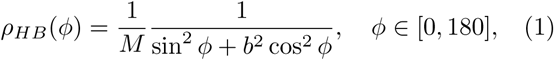

where *M* = 180*/b* is the normalization constant. This result gives a surprisingly good agreement with the experiment (Fig. 1C), especially considering that this model does not take into account the underlying biological processes, e.g. microtubule dynamics. Therefore, while a detailed mathematical model is required to understand how various biological processes control microtubule alignment, the *hairyball* angle distribution Eq.**1** provides a valuable shortcut for the analysis of biological data and parameterizations of microtubule angle distribution.

**FIG. 1.**
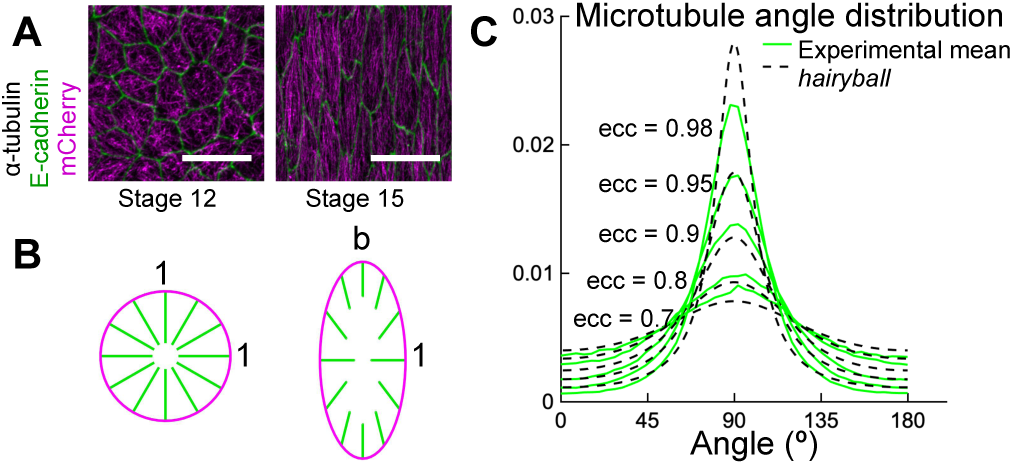
(A) *Drosophila* epidermal cells elongate between the stages of embryonic development 12 and 15, during which the microtubules become more aligned. The bar is 10 *µm*. (B) Stretching a circular cell by a factor *b >* 1 deforms the initially uniform microtubule angle distribution into the *hairyball* distribution, Eq.**1**. (C) The experimental microtubule angle distribution (*green*) and the *hairyball* (*dashed black*) are in good agreement up to eccentricity 0.95. For each eccentricity, the displayed experimental distribution is the mean distribution averaged across cells with the set eccentricity (±0.025 for *ecc* = 0.7 – 0.95 and ±0.005 for *ecc* = 0.98), and is produced as described in Appendix A. The number of cells per eccentricity ranged from 348 to 2748.

### 2. Stochastic simulations demonstrate robustness of microtubule self-organization for a wide range of parameter values

We now explore *in silico* how the microtubule self-organization depends on the parameterization of individual microtubule dynamics and their interactions. To this end we use the same model set-up as previously published [17], as this stochastic model recapitulated microtubule self-organization observed *in vivo* across all range of cell eccentricities. To focus the study on the role of microtubule dynamic instability and interactions and reduce the number of free parameters, we kept the density of the microtubule minus-ends on the cell boundary uniform. The simulations were run on fixed cells, since *in vivo* the cell shape evolve on the longer time-scale (hours), comparing to the time required for the microtubule network statistics to stabilize (several minutes) [17]. We chose the cell shape to be ellipses since we demonstrate below that the averaged experimental cell shape is an ellipse. Finally, the eccentricity range in the simulations 0.7-0.98 mimicked the experimental one.

To capture the dynamic instability, we model microtubules as follows (Fig. 2A). Since the microtubule width (24 *nm*) is much smaller than the typical cell size (2-10 *µm*) [14], we model microtubules as 1d filaments. They are composed of equal length segments, representing microtubule dimers, whose dynamics is governed by a continuous time Markov chain (Fig. 2A, [17, 24]). The microtubule grows (*polymerizes*) at the rate *α*, and shrinks (*depolymerizes*) at the rate *β > α* it switches from the polymerizing to depolymerizing state at the *catastrophe* rate *β′*; and undergoes the reverse switch at the *rescue* rate α′. These rates apart from *β* depend on the concentration of free tubulin dimers in cytoplasm [25], which is reported to vary between 30-75% of the total tubulin in cells *in vivo* [26–28]. However, after the microtubule network stabilizes, the total amount of tubulin in microtubules, and therefore in the cytoplasm, remains approximately constant, leading to approximately constant dynamic instability rates. Since we investigate the statistics of the microtubule network in steady state, we use constant microtubule dynamic instability rates throughout the simulation and set the same rescue rate for a completely depolymerized microtubule as for any non-zero length microtubule. We also exclude the initial period of the simulation from the statistical analysis, until the network reaches steady state (see Appendix C).To account for potentially different microtubule network organizations due to the initial tubulin concentration, we investigate a broad range of parameters. As discussed below, we find that the microtubule angle distribution of the stabilized microtubule network does not sensitively depend on the parameters of the dynamic instability.

**FIG. 2.**
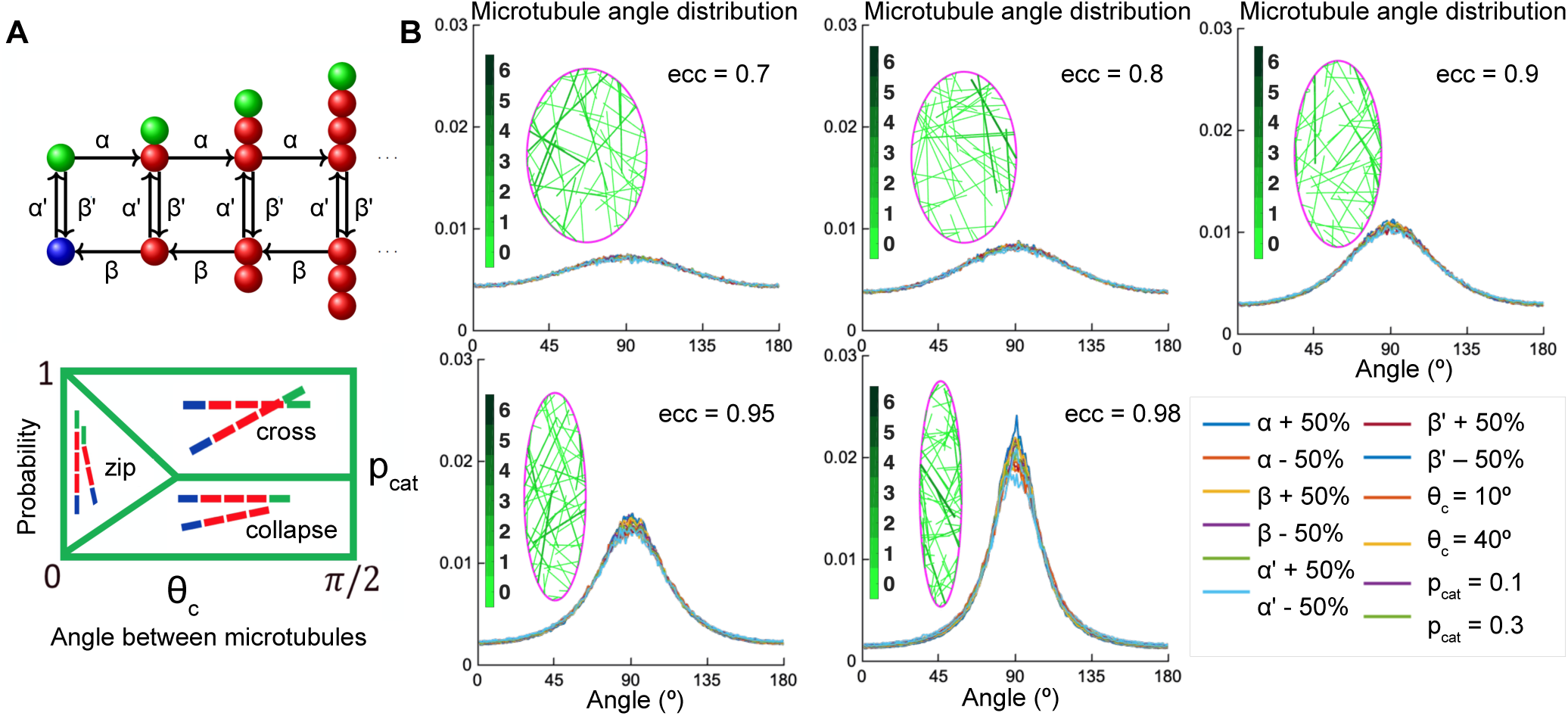
(A, top): Markov chain: each microtubule grows from the minus-end (*blue*) at the rescue rate *α′*, polymerizes at the rate *α* if it is stable with a T.GTP cap (*green*), undergoes catastrophe losing the T.GTP cap at the rate *β′*, depolymerizes at the rate *β*, and regains the T.GTP cap with the rescue rate *α′*. (A, bottom): the angle-dependent probabilities of microtubule interaction scenarios with the critical angle of zipping *θ*_*c*_ and the probability of catastrophe *p*_*cat*_. (B) Sensitivity analysis of the microtubule angle distribution. Left: Snapshots of zipping simulations in cells (*magenta*) of different eccentricities; for the base-level parameters (*α, β, α′, β′*) = (1000, 3500, 4, 1), *θ*_*c*_ = 30 and *p*_*cat*_ = 0.01. Interacting microtubules form bundles, the colorbar indicates the number of microtubules in a bundle. Right: The microtubule angle distributions do not vary significantly in a wide parameter range, suggesting robustness of the microtubule self-organization. The distributions are shown for the variations of(*α, β, α′, β′*) as compared to (1000, 3500, 4, 1), *θ*_*c*_ angle of zipping, and different values of the probability of catastrophe *p*_*cat*_.

We parameterize the angle-dependent microtubule interaction with other microtubules and cell boundaries based on the known interactions in plants [19] and *Drosophila* cells (Fig. 2B, [17, 29]) using two parameters: the critical angle *θ*_*c*_ and the probability of catastrophe *p*_*cat*_. Upon encountering another microtubule at an angle *θ*, if *θ ≥ θ*_*c*_, the growing microtubule undergoes a catastrophe with probability *p*_*cat*_ or crosses the microtubule otherwise. If *θ ≤ θ*_*c*_, it collapses with probability 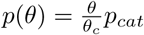, crosses with probability 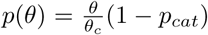, and otherwise bends to change its direction and continues to grow parallel to the existing microtubule (the microtubule is said to *zip*). Upon reaching the cell boundary at an angle *θ ≤ θ*_*c*_, the microtubule zips along it with probability 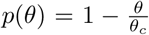, and depolymerizes otherwise [17]. This setup allows us to investigate a broad range of biologically relevant microtubule dynamics scenarios: in an organism, the dynamic instability parameters (*α,β,α*′,*β′*) are linked to the expression of plus-and minus-end binding proteins and severing factors, while the interaction parameters *θ*_*c*_ and *p*_*cat*_ are linked to the presence of crosslinking and motor proteins and temperature-dependent microtubule rigidity [9]. Here we link the model parameters to their dimensional equivalents. The typical observed microtubule growth speed is 0.15 *µm/sec* [17]. Expressing it as *α × d × R*, where *α* = 1000 is the non-dimensional base growth rate, *d* = 8.2 *nm* is the height of one dimer, and *R* is the dimensionality coefficient, we find *R* to be 0.0183 *sec*^−1^. Therefore, the dimensional rates are: the microtubule growth speed *α*_*dim,speed*_ = 0.15 *µm/sec*; shrinking speed *β*_*dim,speed*_ = 0.52 *µm/sec*; rescue rate 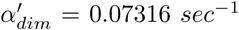; and catastrophe rate 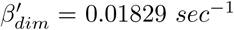.

We varied the parameters (*α, β, α′, β′*) independently, each from 0.5 to 1.5 times the base-line value (*α, β, α′, β′*) = (1000, 3500, 4, 1). We therefore tested the range of dimensional parameters: the microtubule growth speed *α*_*dim,speed*_ = 0.075 0.225 *µm/sec*; shrinking speed *β*_*dim,speed*_ = 0.26 0.78 *µm/sec*; rescue rate 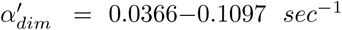; and catastrophe rate 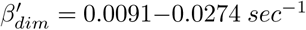. We were representing sample points in the biologically relevant range, since *in vivo* the parameters of microtubule dynamics widely depend on the cell type, with the reported ranges of growth being 0.05 0.5 *µm/sec*; shrinking 0.13 0.6 *µm/sec*; rescue 0.01 0.17 *sec*^−1^; and catastrophe 0.003 0.08 *sec*^−1^ [14, 30, 31].

The simulations (Fig. 2B) gave an unexpected result, that in this large parameter range, the microtubule angle distribution varied only slightly, suggesting that in this model the microtubule self-organization is robust and does not depend on the details of dynamical instability and microtubule interaction parameterization. This suggests that the *in vivo* system could be similarly robust with respect to variations in internal and external environment.

### 3. In vivo manipulations of microtubule dynamics and stability alter microtubule density but not alignment

To test *in vivo* the robustness of microtubule selforganization predicted by the model, we examined how changes in microtubule dynamics and stability affect organization of subapical microtubules in cells of the *Drosophila* embryonic epidermis, where microtubules are constrained to the thin 1 *µm* apical layer of the cell and grow in the plane of the adhesion belt [17]. Using genetic manipulations we could either increase the catastrophe rate *β′*, simultaneously increase the catastrophe rate *β′* and shrinkage rate *β*, or reduce the number of minus-ends, therefore reducing density of the network and thus encounters and zipping between microtubules. In particular, we increased *β′* by overexpressing a dominant-negative variant of End Binding protein 1 (EB1-DN), a form which increases the number of catastrophes without changing other parameters of microtubule dynamics [14, 30]; or increased both *β′* and *β* by overexpressing the protein Spastin, which severs and disassembles microtubules [14, 32, 33]. These proteins were overexpressed using the UAS-Gal4 system, in which the Gal4 protein expressed from a tissue-specific promoter induces over-expression of the protein of interest by binding the Upstream Activating Sequence (UAS) [34]. Specifically, we used *engrailed*::Gal4, which drives expression in stripes along the dorso-ventral axis of embryos, which correspond to posterior halves of each segment ((Fig. 3A)). In this instance we avoided abolishing all microtubules by using mild Spastin overexpression (Fig. 3B). Overexpression of a CD8-Cherry protein, which does not alter microtubules, was used as a control. Finally, we reduced the number of minus-ends using null mutation of the minus-end capping protein Patronin [35], one of the best characterized proteins that protects microtubule minus-ends [36]. Some Patronin protein is present even in homozygous mutant embryos due to maternal contribution, therefore subapical microtubules are reduced but not abolished as shown below.

**FIG. 3.**
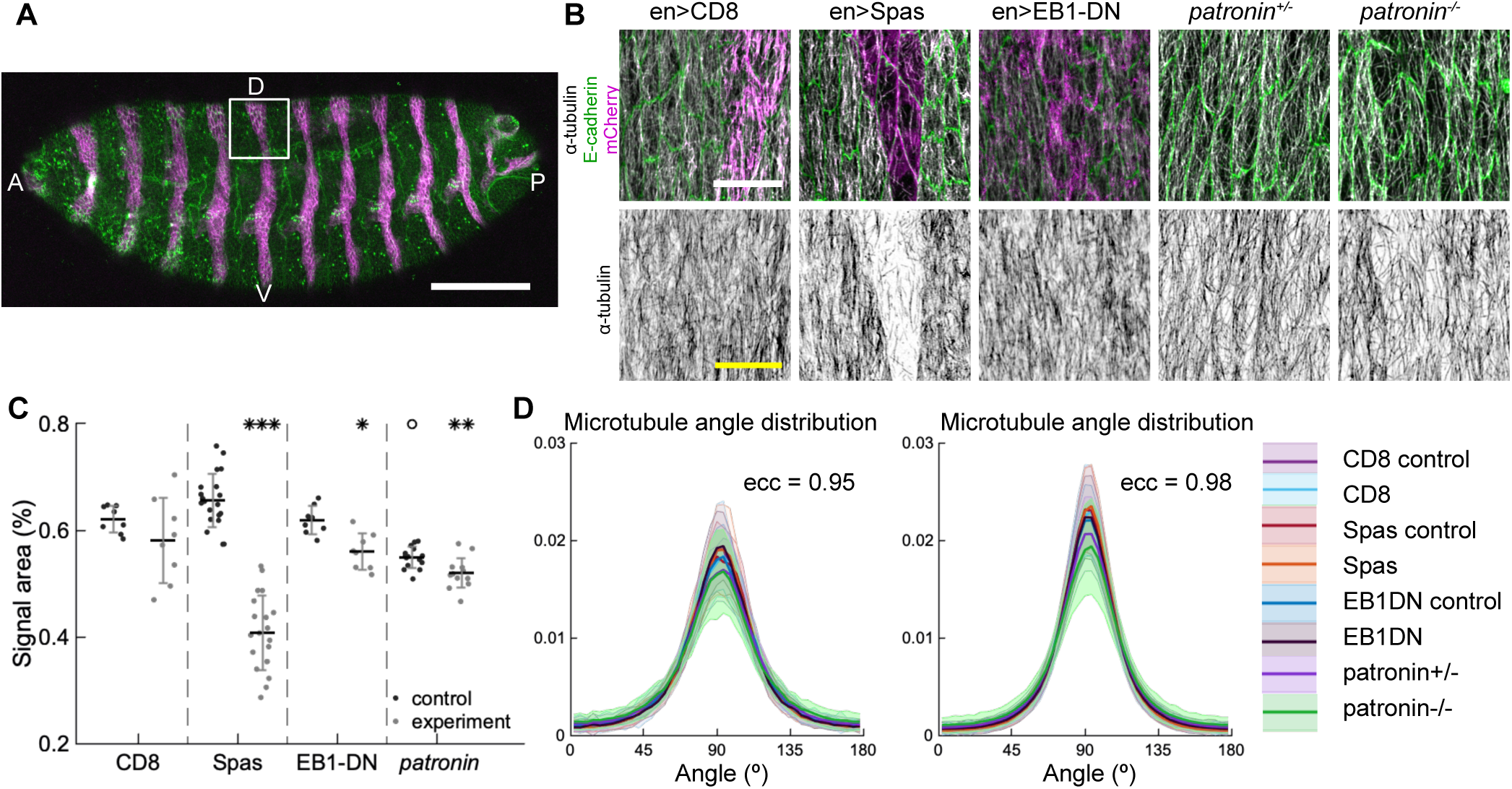
Changes to microtubule dynamics and stability also do not affect their alignment when strong *engrailed*::Gal4 driver is used. (A) An overview image of a *Drosophila* embryo at the stage 15 of embryonic development with cell outlines visualized with E-cadherin (green), and *engrailed*-expressing stripes are visualized by direct fluorescence of mCherry (magenta). The white square demonstrates the areas used for analysis on microtubule organization as shown in (B). Anterior (A), posterior (P), dorsal (D), and ventral (V) sides of embryos are labelled. Scale bar - 100 *µm*. (B) Apical view of epidermis from control embryos and with altered microtubules. Left-to-right: embryos with CD8-Cherry (control), Spastin (Spas), and EB1-DN expressed using *engrailed*::Gal4, heterozygous *Patronin*^-/+^, and homozygous *Patronin*^-/-^ embryos. Cells expressing CD8-Cherry and EB1-DN are visualized by direct fluorescence of mCherry directly fused to respective proteins, whereas cells expressing Spastin are visualized by coexpression of CD8-Cherry (*magenta*, top row). Cell outlines were visualized by immunostaining against E-cadherin (*green*, top row), and microtubules by immunostaining against *α*-Tubulin (*white*, top row; *black*, bottom row). Scale bar - 10 *µm*. (C) Quantification of microtubule density in each genotype. Internal controls (cells not expressing *paired*::Gal4) were used for CD8-Cherry, Spastin, and EB1-DN overexpression. For *Patronin*, heterozygous and homozygous embryos were compared. *** - *p* < 0.0001, ** - *p* < 0.001, * - *p* < 0.01 in comparison to respective control; *°* - *p* < 0.01 in comparison to CD8-Cherry control. (D) The microtubule angle distributions for each eccentricity (*±*0.025 for *ecc* = 0.95 and *±* 0.005 for *ecc* = 0.98) do not significantly differ between all genotypes and relatively to controls. The distributions are shown as mean (solid line) with standard deviation (shading). The number of cells per eccentricity per genotype ranged from 32 to 833.

We quantified how the above manipulations altered organization of the microtubule network in cells by obtaining images of microtubules stained with an antibody recognizing *α*-Tubulin, and analyzing them on a cell-by-cell basis. From these images we obtained two types of information about microtubule organization, namely about density and alignment of the network (see Appendix F).First, we determined how the manipulations altered the amount of microtubules in cells, by quantifying the percent of the cell area covered by *α*-Tubulin signal as a proxy for the number of microtubules in cells. Second, we determined if the genetic manipulations altered the microtubule alignment. To do this we determined the direction and magnitude of change of the *α*-Tubulin signal at each position within the cell (see Appendix G and [17]), and produced the microtubule direction distributions.

Amounts of protein expressed using UAS-Gal4 system increases with developmental time, whereas when using mutant embryos, amounts of protein supplied from mothers in eggs, the maternal contribution, decreases with time as they cannot produce their own protein. Therefore, we focused on the late developmental stage of *Drosophila* embryo development (stage 15). The over-expression of both Spastin and EB1-DN reduced the area of *α-*Tubulin signal in cells (*p*-values *p* < 0.0001 and *p* = 0.003, respectively, Fig. 3B-C), consistent with their functions. Similarly, the area of *α*-Tubulin signal was reduced in heterozygous *Patronin*^+/-^ embryos in comparison to wild-type controls (*p*-value *p* = 0.02, Fig. 3A-B), and even further reduced in homozygous *Patronin*^-/-^ embryos (*p*-values *p* = 0.0003 and *p* = 0.002 in comparison to wild-type control and heterozygous siblings, respectively, Fig. 3B-C). This result is consistent with the dose-dependent protection of the microtubule minus-ends by Patronin. Despite these large changes in microtubule density, the microtubule angle distributions did not significantly differ between all genotypes and relatively to controls in this case (Fig. 3D), which supports robustness of microtubule self-organization despite different microtubule dynamics and amount.

To capture a wide range of eccentricities we used the embryos at different stages of development when the epidermal cells progressively elongate from eccentricities around 0.7 to 0.98 (stages 12 through 15). To this end we used *paired*∷Gal4 to overexpress Spastin, EB1-DN, and CD8-Cherry, which although leads to milder over-expression than *engrailed*∷Gal4, is expressed in broader stripes along the dorso-ventral axis of embryos. Overexpression of Spastin reduced *α*-Tubulin signal area in comparison to both neighboring cells, which did not express *paired*∷Gal4, and control cells expressing CD8-Cherry (both *p*-values *p* < 0.0001, Figure blue S1A-B). Similarly, the *α*-Tubulin signal area was reduced in embryos homozygous for *Patronin*^-/-^ in comparison to heterozygous siblings (*p*-value, *p* = 0.04, Figure blue S1A-B). We suggest that no difference between heterozygous *Patronin*^+/-^ and wild type control observed here, might be due to broader developmental time, which makes more difficult to detect changes. Overexpression of EB1-DN did not change the area covered by the *α*-Tubulin signal per cell (*p*-value, *p* = 0.98, Figure blue S1A-B). This can be explained by either weaker expression of *paired*∷Gal4 in comparison to *engrailed*∷Gal4, or lower amounts of Gal4 at earlier developmental stages [14]. The microtubule angle distributions for each eccentricity (binned at a particular eccentricity ± 0.025) did not significantly differ between all genotypes and relatively to controls (Figure blue S1C). Although only two of the above manipulations affected microtubule density, these results further support that microtubule self-organization is indeed robust *in vivo*.

### 4. An analytical model shows that microtubule self-organization depends on the cell geometry and minus-end distribution

Given that the microtubule angle distribution *in silico* and *in vivo* only weakly depends on microtubule interactions, we propose a minimal mathematical model with non-interacting microtubules, which is analytically tractable. Here, independent microtubules cross upon reaching another one, and fully depolymerize upon reaching a cell boundary. Their averaged behavior is the average of 1d behaviors of individual microtubules growing from different positions on the cell boundary.

Our model setup is as follows (Fig. 4A). Consider a convex 2d cell with the boundary parameterized by the arclength-coordinate *ζ* increasing in a counter-clockwise direction. From microtubule minus-ends distributed with density *ρ*(*ζ*) on the cell boundary, the microtubules grow into the cell interior at an angle *θ* to the cell boundary, making the angle *ϕ* with respect to the horizontal. Note that the cell shape is fully determined by the function *a*(*ζ, θ*) – the length of a cross-section that starts at *ζ* at an angle *θ* with respect to the boundary. When *a*(*ζ, θ*) is considered as a function of(*ζ, ϕ*), we denote it by *ã* (*ζ, ϕ*) to avoid confusion. Microtubules undergo dynamic instability by switching between the states of growth, shrinking, catastrophe and rescue at the rates *α, β, β*′, and *α*′ respectively. After fully depolymerizing, they regenerate at the rate *α*′ from the same minus-end but in a new direction at an angle *θ* taken from a uniform distribution on [0, 180]. Thus, on a large fixed time interval *t* ∈ [0,*T*] a microtubule undergoes a large number *N* of growth and shrinkage lifetimes separated by periods of average duration 1*/α*′ when it has zero length.

**FIG. 4.**
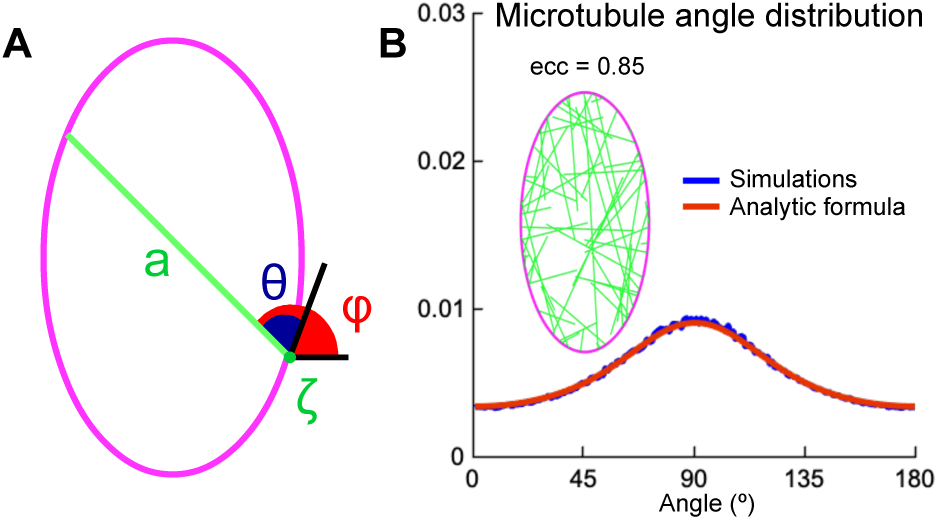
(A) Analytical model setup: cell shape is parameterized by the arclength *ζ* along the cell boundary. At the minus-end *ζ* a straight microtubule grows at an angle *θ* (or *ϕ*) with respect to the cell boundary (or horizontal); its maximum length is the cross-section *a* of the cell. (B) Left: snapshot the simulations with non-zipping independent microtubules for cells of eccentricity 0.85. Right: agreement between the microtubule angle distributions Eq.**9** (*red*) and the stochastic simulations (*blue*).

The first quantity of interest is the microtubule mean survival time. Since in the *in vivo* and *in silico* the microtubule angle distribution is length-weighted, we include the general case of weighting the mean survival time by a function *γ*(*x*) of microtubule length *x*. Then the mean survival times *f*(*x*) and *g*(*x*) of polymerizing and depolymerizing microtubules of length *x* satisfy

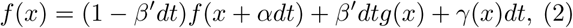

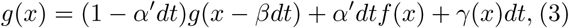

where the terms on the right-hand side are the contributions from growing (and shrinking in the *g* case), switching, and the time increment weighted by *γ*(*x*), which we specify below. Here *dt* is a small time-increment. Expanding Eq.**2-3** in Taylor series and neglecting terms of the second and higher order in *dt*, we obtain that *f*(*x*) and *g*(*x*) are governed by

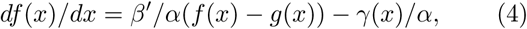

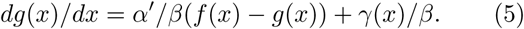

Their difference *h*(*x*) = *f*(*x*) *− g*(*x*) satisfies

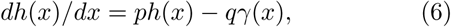

where 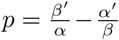 and 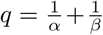. We assume that *g*(*a*) = 0, i.e. once the microtubule reaches the cell boundary at the length *a*(*ζ, θ*), it quickly depolymerizes. Finally, for a zero-length microtubule *g*(0) = 0, and hence *h*(0) = *f*(0). Note that the only quantity of interest is *f*(0) = *h*(0), since it is the lifetime of a microtubule when it starts in a growing state with zero length. The two choices *γ*(*x*) = 1 and *γ*(*x*) = *x* give the solutions for the not-weighted and the length-weighted mean survival times, denoted *f*_*nw*_(*x*) and *f*_*w*_(*x*) respectively

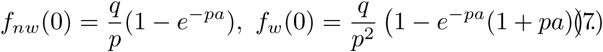

For each microtubule minus-end location *ζ*, the average time between any two re-growths of a microtubule is the sum of the averaged waiting time 1*/α′* and the average of the non-weighted lifetime over all the growth angles *θ*

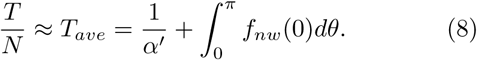

Then the average number of lifetimes with direction (*ϕ, ϕ* + δ*ϕ*) with respect to the horizontal is *Nδϕ/ π*, and their contributions to the length-integral is *f*_*w*_(0)*Nδϕ/π* = (*f*_*w*_(0)*/T*_*ave*_)*T/ πδϕ*. Integrating it over the cell boundary weighted by the density of minus-ends *ρ*_*m*_(*ζ*) and using Eq.**8**, we obtain the length-weighted microtubule angle distribution

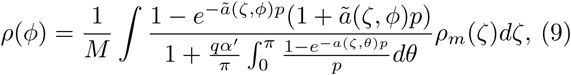

where *M* is the normalization constant. The cell cross-section is denoted by either *a*(*ζ, θ*) or ã (*ζ, ϕ*) depending on its arguments. This analytical prediction matches the stochastic simulations (Fig. 4B).

The dependency of the resulting microtubule angle distribution on the biological rates reduces to two parameters

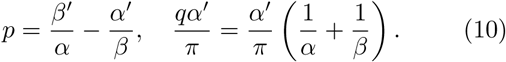

If both are small, *|p| ≪* 1 and 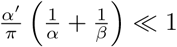, as is always observed in biological systems (see above), the microtubule angle distribution formula can be significantly simplified. In particular, for small *p* the distribution Eq.**9** reduces to

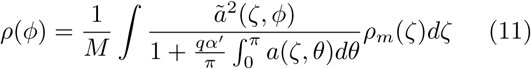

to leading order, and to

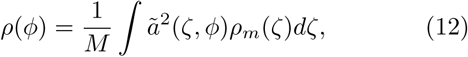

when, in addition, 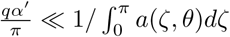. This becomes exact in the limit of deterministic microtubules (*β* = ∞, *β′* = 0). Note that while *p* is required to be non-negative in models of microtubules on an infinite line [24], our setup does not have this restriction, as the microtubule lifetimes *f*_*w*_(0) and *f*_*nw*_(0) are positive even for negative *p*. Furthermore, the analytical microtubule angle distribution Eq.**9** is independent of the multiplicative change in the minus-end density *ρ*_*m*_(*ζ*), which would be absorbed into the normalization constant *M*. Only non-trivial changes to the density of minus-ends that vary along the cell boundary affect the microtubule angle distribution.

It is required that *α′ ≪ α* and *α′ ≪ β* for the second parameter in Eq.**10** to be small, and *β′ ≪ α* for the first one to be small as well. Therefore, the microtubule angle distribution depends only on the cell geometry and the minus-end distribution, and the underlying reason for that is the separation of time-scales in microtubule dynamics: the rates of polymerization and depolymerization are much higher than those of catastrophe and rescue, which is always observed *in vivo*.

### 5. The analytical model accurately predicts microtubule-self organization using experimental cell shape data and distribution of microtubule minus-ends

In order to validate the efficacy of our analytical model in prediction of microtubule organization, we first determined the localization of microtubule minus-ends. Since Patronin localizes at the microtubule minus-ends, we analyzed its distribution in epithelial cells in the *Drosophila* embryonic epidermis using Patronin-YFP [35]. As expected, Patronin-YFP is mostly localized at the cell boundaries with few speckles inside cells (Fig. 5A). We quantified the distribution of Patronin-YFP at the cell boundaries by measuring its asymmetry, namely the ratio of Patronin-YFP average intensity at dorso-ventral borders to that of anterior-posterior borders (see Appenxis H, [37], and Fig. 5B). The asymmetry of Patronin distribution *B*(*ecc*) was a linear function of the cell eccentricity (Fig. 5C), suggesting that Patronin becomes enriched at the dorso-ventral boundaries as the embryo develops and cells elongate. Additionally, when comparing boundaries in cells with similar eccentricities (Stage 15 embryos only), the intensity of Patronin-YFP was decreasing with the border angle relatively to anterior-posterior axis of the embryo (Fig. 5D). Several lines of evidence support that the observed enrichment of Patronin-YFP at the dorso-ventral boundaries is due to the asymmetry of cell-cell adhesion in these cells. Indeed, microtubule minus-ends were shown to be tethered by cell-cell adhesion in some epithelial cells [38]. At the same time, E-cadherin, the key component of cell-cell adhesion in *Drosophila* embryonic epidermis, is asymmetrically distributed in stage 15 embryos [14], with an enrichment at the dorsal-ventral borders similar to that of Patronin. Finally, asymmetries of both Patronin and E-cadherin progressively increase from stage 12 to 15 of *Drosophila* embryonic development (Fig. 5C and N.A.B. personal communication).

**FIG. 5.**
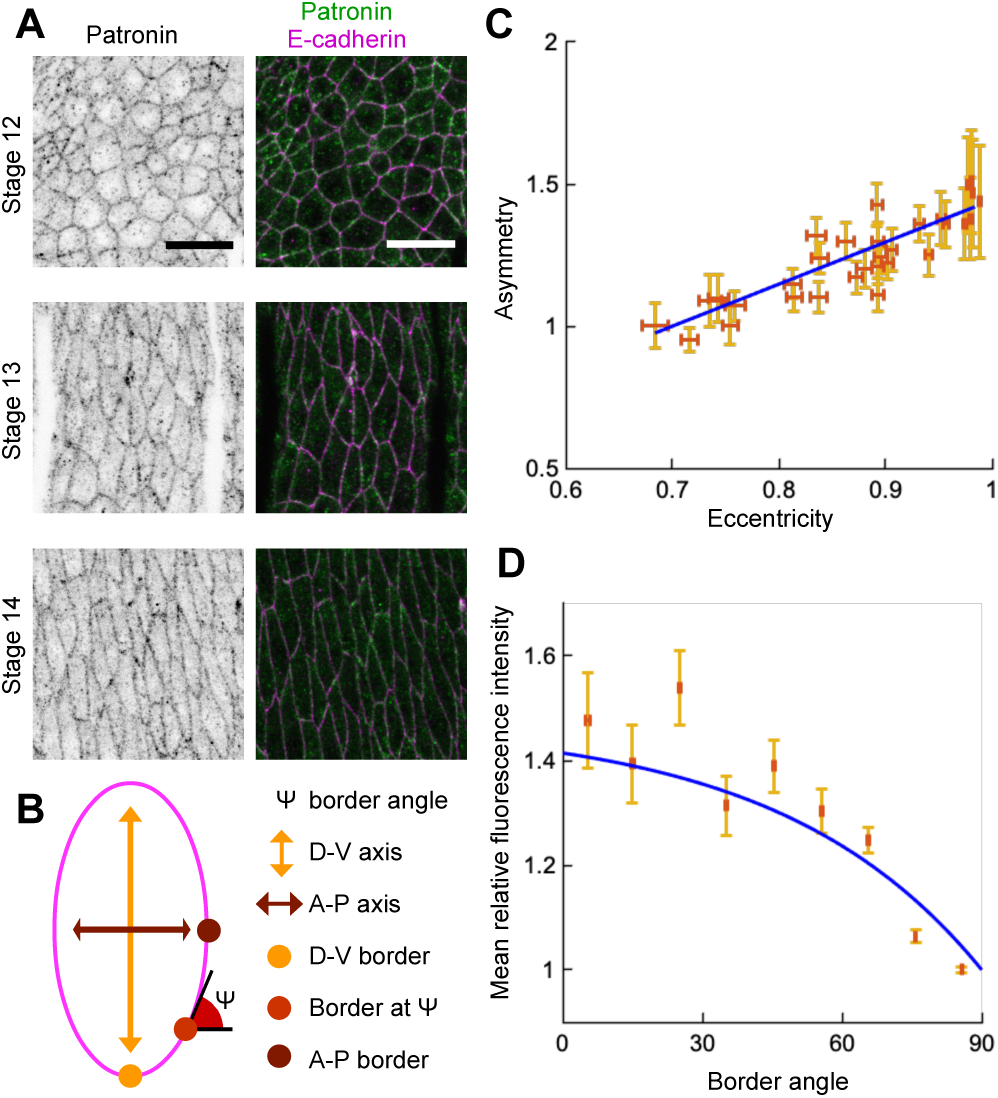
Localization of Patronin-YFP in the *Drosophila* embryonic epidermis. (A) Apical view of embryonic epidermis at stage 12 (top), 13 (middle), and 15 (bottom), visualized with Patronin-YFP (*grey*, left; *green*, right), and E-cadherin immuno-staining (*magenta*, right). Scale bar, 10 *µm*. (B) Schematic of a cell. (C) Asymmetry of Patronin-YFP localization increases linearly with eccentricity. Each dot represents average values of Patronin-YFP asymmetry and cell eccentricity in a single embryo. Error bars are SD. Solid line visualizes the linear fit of the form *B*(*ecc*) = 1 + *C*_1_(*ecc* − 0.7)*/*(0.98 − 0.7), *C*_1_ = 0.4144. (D) Mean relative amounts of Patronin-YFP as a function of the border angle *ψ* in stage 15 (eccentricity 0.98) *Drosophila* embryonic epidermis, normalized by its value at the vertical long sides (0-10*°*). Borders were binned at 10*°* intervals relatively to embryonic anterior-posterior axis, and intensity was averaged for each bin (mean *±* SD). The solid line represents the exponential fit for the intensity as 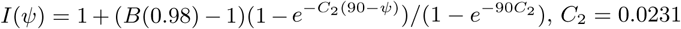, *C*_2_ = 0.0231.

To use this in our analytical model, we used least-squares to simultaneously fit the asymmetry data with a linear function of eccentricity, and the normalized intensity of Patronin-YFP with an exponential function of the cell border angle. We imposed a constraint that the asymmetry value at eccentricity 0.98 (Stage 15) is the same as the normalized intensity of Patronin-YFP at the dorso-ventral border (border angle 0*°*). The resulting formula used in the analytical model is

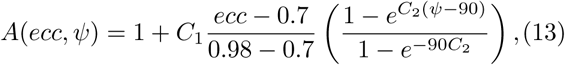

where *ψ* is the cell border angle with respect to the horizontal, *C*_1_ = 0.4144 and *C*_2_ = 0.0231.

Next, we determined the average cell shape. In the tissue, each cell has a unique shape, and cells with the same eccentricities may differ significantly in their geometry. Therefore, to test and validate the analytical solution of microtubule self-organization we have generated masks of epithelial cells in the *Drosophila* embryonic epidermis, which provided us with coordinates of cell boundaries (see Appendix G).Dividing all cell shapes into groups by eccentricity, we computed the average cell shape for each group as follows. First, the cells were re-centered to have their centers of mass at the origin. We then rotated them so that they are elongated along the vertical axis (the direction of elongation is the first singular vector, see Appendix G). Finally, we rescaled all the cells to have unit area. The average of distance from the center of mass in a particular direction to the cell boundary traced the boundary of the averaged cell. Surprisingly, we found that the average cell shape for a given cell eccentricity is an ellipse (Fig. 6).

**FIG. 6.**
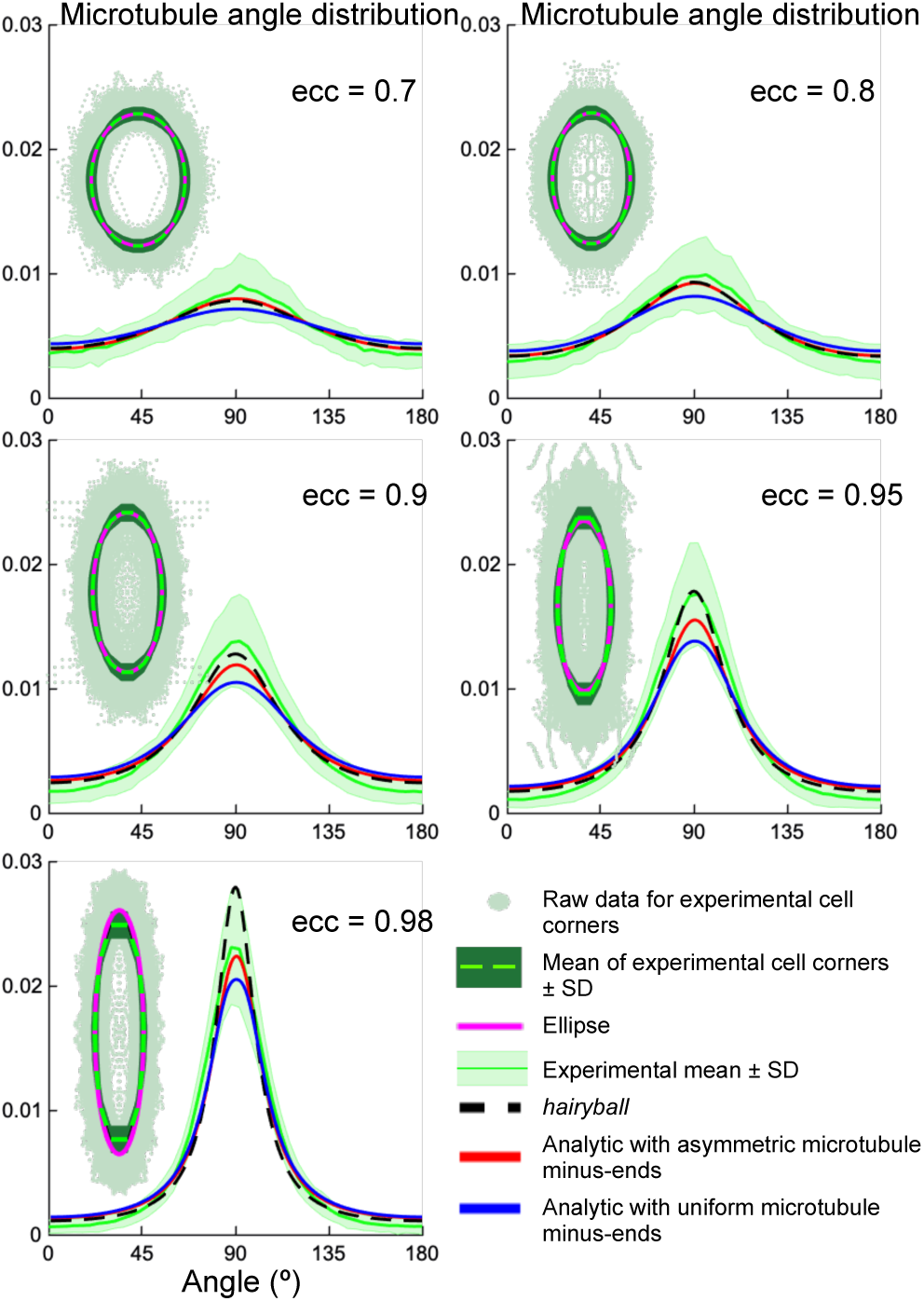
Top left of each graph: experimental cell shapes. The experimental cell boundary data points (*light green points*), its standard deviation (*darker green envelope*) around the radial mean (*dark green line*). The ellipse (*dashed magenta*) closely approximates the experimental mean shape. The graphs show the corresponding microtubule angle distributions. The analytical distribution with asymmetric minus-ends (*red*) has better agreement with the experimental mean (*dark green line*) (±*SD* is the *light-green envelope*), comparing to the uniform minus-end density (*blue*); the *hairyball* has good agreement up to eccentricity 0.95 with the experimental mean.

The analytical microtubule angle distribution Eq.**9** computed on an averaged cell shape using the experimental minus-end density Eq.**13** gives surprising agreement with experimental *in vivo* data (Fig. 6). This agreement is better for the case of asymmetric minus-end distribution, comparing to the uniform one, which supports our prediction that the minus-end distribution does indeed influence the microtubule self-organization.

## III. DISCUSSION

Here we present several novel findings describing the fundamental rules underlying self-organization of microtubule networks in epithelial cells. Firstly, we have shown robustness of microtubule self-organization both *in silico* and *in vivo*; secondly, in addition to the known importance of cell shape to microtubule organization [17], our minimal analytical model predicted the importance of the asymmetric minus-end distribution as the only other impactful parameter, which we then confirmed *in vivo* in *Drosophila* epidermal cell; and finally, that the origin of robustness is the intrinsic separation of time-scales of microtubule dynamic instability.

### A. Robustness

Biologically, the discovered robustness of microtubule organization makes perfect sense, given the fundamental importance of microtubule functions in a cell. Most of the intracellular trafficking events require microtubules for the delivery of various cellular components to their relevant biological locations by motor proteins [39, 40]. This process must be reliable, as mislocalization of cellular components leads to cell death or disease [41–43]. However, the delivery mechanism is highly stochastic, given the nature of the microtubule dynamic instability and the dynamics of molecular motors [44–46]. We suggest that the overall organization of the microtubule network is likely to guide the net outcomes of intracellular trafficking while minimizing the impacts of stochastic effects. Indeed, our mathematical model shows that while the dynamics is stochastic, the dependence of the microtubule self-organization only on the slowly evolving parameters, such as the cell shape and the density of the minus-ends on the cell boundary, makes the average behavior of the system deterministic on long time-scales. Averaging the cell shape over the cells of the same eccentricity brings us to the tissue level: while most of the cells of given eccentricity in biological tissues are polygons, we found that the average cell shape of a particular eccentricity in the tissue is an ellipse. This suggests that though there might be differences in microtubule organization in individual cells at a given point in time, on the tissue scale the average shape and hence microtubule organization is robust.

One of the findings of our analytical model is that the robustness of microtubule self-organization exists only as long as the microtubule dynamics exhibits a separation of time-scales, *α*′, *β*′ ≪ *α, β*, a rule which is observed in all published data about microtubule dynamic parameters [14, 31, 47–53] Our mathematical model shows that if this rule is not observed, the microtubule organization becomes sensitive to changes in these rates. As these rates depend on multiple internal and external factors, such as stochastic changes in gene expression and environment temperature, the microtubule organization would be unpredictable at any certain time in a cell in a biological tissue. Therefore, breaking this rule will impair cellular function over time, which suggests that any mutations that led to such change were likely to cause cell lethality and did not fix in evolution.

### B. *Hairyball* distribution

We demonstrated that the microtubule angle distribution is accurately predicted by the *hairyball* distribution. This excellent agreement with the experimental data remains an open question, as we were unable to show that *hairyball* has any relation to the analytical distribution, neither as an approximation nor as a limiting case. We suggest that its best use is as a simple ad-hoc formula to parameterize the microtubule angle distribution in cells up to eccentricity 0.95 in investigating, for example, correlations and interdependencies between the microtubule network organization (its statistical properties, e.g. the overall direction and spread) and dynamic intracellular processes, e.g. signaling and transport.

### C. The importance of microtubule-microtubule interactions

While we show that the microtubule self-organization does not depend on the interaction of microtubules, we admit that this is true for the measure of self-organization being the time-averaged microtubule angle distribution. When using this measure, such effects of microtubule-microtubule interactions as zipping and bundling disappear due to the time-averaging of the dynamics. From the biological point of view, what matters for the organism is the long-term behavior, because most of the processes such as microtubule-based transport occur at much longer time scales than microtubule network rearrangements [14, 54, 55]. Therefore, while bundling and zipping are aspects of the short-term behavior, we hypothesize that it is the averaged microtubule angle distribution, which affects the tissue behavior on the long time-scale, with the long term behavior being more important than snapshots. However, we hypothesize, that such effects as bundling and zipping will affect short-term intracellular transport. For example, the presence of Spastin, the microtubule severing protein, leads to a change in the delivery of the E-cadherin, the protein responsible for the cell-cell adhesion which is delivered along the microtubule network [14]. A more detailed modeling approach that includes the effect of microtubule-microtubule interaction on intracellular transport is outside the scope of this article, and will be considered in a separate publication.

### D. Our system is a particular but generalizable scenario

We suggest that our findings are applicable beyond apical microtubules in the *Drosophila* embryonic epidermis, the dynamics of which is quasi-2d. Previously we have demonstrated that a similar relationship between microtubule organization and cell shape is observed in other *Drosophila* epithelia, including cells in pupal wings and ovaries [17]. These are only two of the examples where our findings are likely to hold true. There are multiple other instances, whereupon maturing and differentiating epithelial cells develop an apical microtubule meshwork, including cells of mammalian airways, and even cells in culture [16, 56, 57]. Furthermore, the same rules are likely to apply to squamous cells, where despite having specialized apical microtubules, the cells’ depth is so small, that microtubules are constrained within a thin plane similar to that in our experimental model [58, 59]. The validation of our findings in other cell types and other evolutionary divergent organisms, as well as how the discovered robustness of microtubule selforganization ensures reliability of intracellular transport are important questions for future research.

## ACKNOWLEDGMENTS

This research is supported by The Maxwell Institute Graduate School in Analysis and its Applications, a Centre for Doctoral Training funded by the UK Engineering and Physical Sciences Research Council grant EP/L016508/01, the Scottish Funding Council, Heriot-Watt University and the University of Edinburgh (A.Z.P.); BBSRC BB/P007503/1 (N.A.B.); Royal Society of Edinburgh and the Scottish Government Personal Research Fellowship (L.C.); and Leverhulme trust grant RPG-2017-249 (to L.C. and N.A.B).

## Author contributions

N.A.B. and L.C. designed the research and experiments; A.Z.P. and L.C. performed stochastic simulations; A.Z.P., A.M.D., and L.C. developed analytical model; N.A.B. did biological experiments; A.Z.P., N.A.B., and L.C. wrote the manuscript. The authors declare no conflict of interest. N.A.B and L.C. contributed equally to this work.

## Appendix A: Computing the eccentricity of a cell

We uniformly distributed points inside the experimental cell boundary data and used the singular value decomposition on the resulting dataset. The eccentricity then is *ecc* = (1 − (*a/b*)^2^)^1*/*2^, where *a* < *b* are the singular values.

## Appendix B: Derivation of the *hairyball* distribution

Upon stretching a circular cell by a factor of *b* > 1 in the vertical into an ellipse with eccentricity *ecc* = (1 − 1*/b*^2^)^1/2^, the angles *θ* with uniform distribution *R*(*θ*) = 1*/*180 are mapped into the angles with the distribution *ρ*(*ψ*) such that *R*(*θ*) = *Jρ*(*ψ*), where *J* is the Jacobian of the stretching map. Since *R*(*θ*) = *const, ρ*(*ψ*) ∝ *J* ^−1^ =(sin^2^(*ψ*) + *b*^2^ cos^2^(*ψ*) ^−1^.

## Appendix C: Parameters of stochastic simulations

Along the cell boundary discretized with 180 points, 200 microtubule minus-ends were placed uniformly. The short axis of the cell was kept constant at 60 numerical dimer lengths. The critical angle *θ*_*c*_ was varied in the range (0, 10, 20, 30, 40) degrees, and the probability of catastrophe *p*_*cat*_ was varied in the range (0.01, 0.1, 0.2, 0.3). We imposed microtubules zipping along a cell boundary undergo catastrophe upon reaching the tip of the ellipse. This mirrors the scenario *in vivo* that due to the high stiffness [60], microtubules undergo catastrophe upon reaching an acute cell corner (Fig. 1A). All the simulations were run to the non-dimensional time *T* = 10, which corresponds to 550 *sec*. At the start of each experiment all the microtubule lengths were set to zero. We binned the microtubule angles with respect to the horizontal in 180 one-degree bins, and averaged these histograms in the last 2.5 non-dimensional time-units, which corresponds to 250 points. The *in silico* model is available from the authors upon request.

## Appendix D: Fly stocks

*paired*∷Gal4, *engrailed*∷Gal4, *UAS*∷CD8-Cherry (Bloomington stock numbers1947, 32, and 27392, respectively), *UAS*∷EB1-DN [14], *UAS*∷Spastin [61], *Patronin*^*05252*^, and Patronin-YFP [35]. The flies and embryos were kept at 18*°C*.

## Appendix E: Embryo fixation and antibody staining

The embryos were fixed as described in [17]. In brief, the staged embryos were dechorionated in 50% bleach for 4 min, and then fixed in 1:1 10% formaldehyde (methanol free, #18814, Polysciences Inc.) in PBS:heptane for 20*min* at room temperature (RT) and post-fixed/devitellinized for 45*s* in 1:1 ice-cold methanol:heptane. Finally, embryos were washed three times in ice-cold methanol, kept in methanol between 6 and 24*h* at − 20*°C*, rehydrated in 1:1 PBS with 0.3% Triton X-100 (T9284, Sigma):methanol, and washed one time in PBS with 0.3% Triton X-100. Rehydrated embryos were blocked for 2*h* in 5% Native Goat Serum (ab7481, Abcam) in PBS with 0.3% Triton X-100. Primary antibody incubations were carried out overnight at 4*°C*. Primary antibodies used were mouse anti-*α*-Tubulin 1:1,000 (T6199, Sigma), and rat anti-E-cadherin 1:50 (DCAD2, Developmental Studies Hybridoma Bank). Incubation with secondary antibody was performed for 2*h* at 25*°C*. Alexa Fluor fluorophore Alexa Fluor 488- and 647-coupled secondary antibodies (Jackson ImmunoResearch) were used in 1:300. Finally, embryos were mounted in Vectashield (Vector Laboratories).

## Appendix F: Image acquisition

All images were acquired at RT (20-22*°C*). For quantification of microtubule self-organization, we acquired 16-bit depth images on the Zeiss AiryScan microscope, using 60x objective lens. A *z*-stack of 8 sections with 23.5 *px/µm* in XY resolution and 0.38 *µm* distance between *z*-sections were taken. All processing was done at 6.5 power in the ZEN software. For an analysis of Patronin-YFP distribution, an upright confocal microscope (FV1000; Olympus) using 60 × 1.42 NA oil PlanApoN objective lens was used. 16-bit depth images were taken at a magnification of 12.8 *px/µm* with 0.38 *µm* between *z*-sections using Olympus Fluoview FV-ASW.

The sub-apical domain of the cell was determined using the E-cadherin junctional signal as a reference for its basal limit, and the absence of *α*-Tubulin signal as a reference for its apical limit. All imaging was done on dorso-lateral epidermal cells, with excluded the leading edge (first row) cells, given its different identity to the rest of the dorso-lateral epidermal cells. Average projections were made using Fiji (www.fiji.sc) for measuring signal area and Patronin-YFP distribution. Figures were assembled using Adobe CS3 Photoshop and Illustrator (www.adobe.com). Processing of images shown in figures involved adjusting of gamma settings.

## Appendix G: Image analysis to quantify microtubule organization

We used the same workflow as described in the [17]. In short, E-cadherin signal from a max intensity projection was used to obtain cell outlines using Packing Analyser V 2.0 software [62]. These cell outlines were used to identify each cell as an individual object and fit it to an ellipse to calculate eccentricity and the direction of the ellipse major axis using a script in MATLAB R2017b (Mathworks, www.mathworks.co.uk). The *α*-Tubulin signal within each cell was filtered using the cell outline as a mask, and the magnitude of the signal according to its direction analyzed by convolving the filtered *α*-Tubulin signal with two 5 × 5 Sobel operators [17]. The resulting magnitude gradient and direction gradient were integrated into a matrix to assign each pixel a direction and magnitude of change in the intensity of *α*-Tubulin signal. To reduce the noise, the pixels with the magnitude less than 22% of the maximum change were discarded. The remaining pixels were binned with respect to their direction gradient (bin size= 4 degrees). The resulting histogram was normalized. The script is available at https://github.com/nbul/Cytoskeleton.

To calculate the area of *α*-Tubulin signal within each cell, a custom-made MATLAB script was used. The average projected images were adjusted such that 0.5% of the data with lowest intensities were set to black and the 0.5% of the data with highest intensities were set to white in order to compare datasets across genotypes and compensate for potential variability of antibody staining and laser power. The threshold was then calculated using Otsu’s method, and a binary image was created using the calculated threshold multiplied by 0.7. This multiplication parameter was determined empirically by testing images from different experiments. Finally, the percent of pixels above threshold was calculated for each cell. The script is available at https://github.com/nbul/Cytoskeleton.

## Appendix H: Image analysis to quantify Patronin-YFP distribution

The average projection of each *z*-stack was done using Fiji software, and segmented using Packing Analyzer v2.0. The resulting binary images with coordinates of cell-cell borders and vertices were used together with unprocessed average projection by a custom-made MATLAB scripts to extract the following values: apical cell area, cell eccentricity, average cell orientation within the image, orientation of individual cell-cell borders relative to average cell orientation, and mean intensity of each individual border. Only the cells that were completely within the image were taken for quantification. Bristle cells were excluded by their size. Only the borders that are between two cells that were completely within the image were quantified, and the borders adjacent to bristle cells were excluded. The background signal was determined by binarizing images with an adaptive threshold, which uses local first-order image statistics around each pixel and is very efficient in detecting puncta. The mean background signal of cells that were completely within the image was subtracted from mean intensities of cell-cell borders. The values for each type of border within single embryos were averaged, and the average intensity of borders with 40-90*°* orientation was divided by the average intensity of borders with 0-10*°* orientation to produce a single value of asymmetry for each embryo. Finally, to produce distribution of signal intensity by angle, borders of all embryos at stage 15 of development were pulled and binned using 10*°* bins. The script is available at https://github.com/nbul/Intensity.

## Appendix I: Statistical analysis

All data was analyzed using Graphpad Prism 6.0c (www.graphpad.com). Samples from independent experiments corresponding to each genotype were pooled and tested for normality with the Shapiro-Wilk test. The *α*-Tubulin signal area were analyzed using ANOVA with post-hoc t-test.

## Supplementary information

**FIG. 1.**
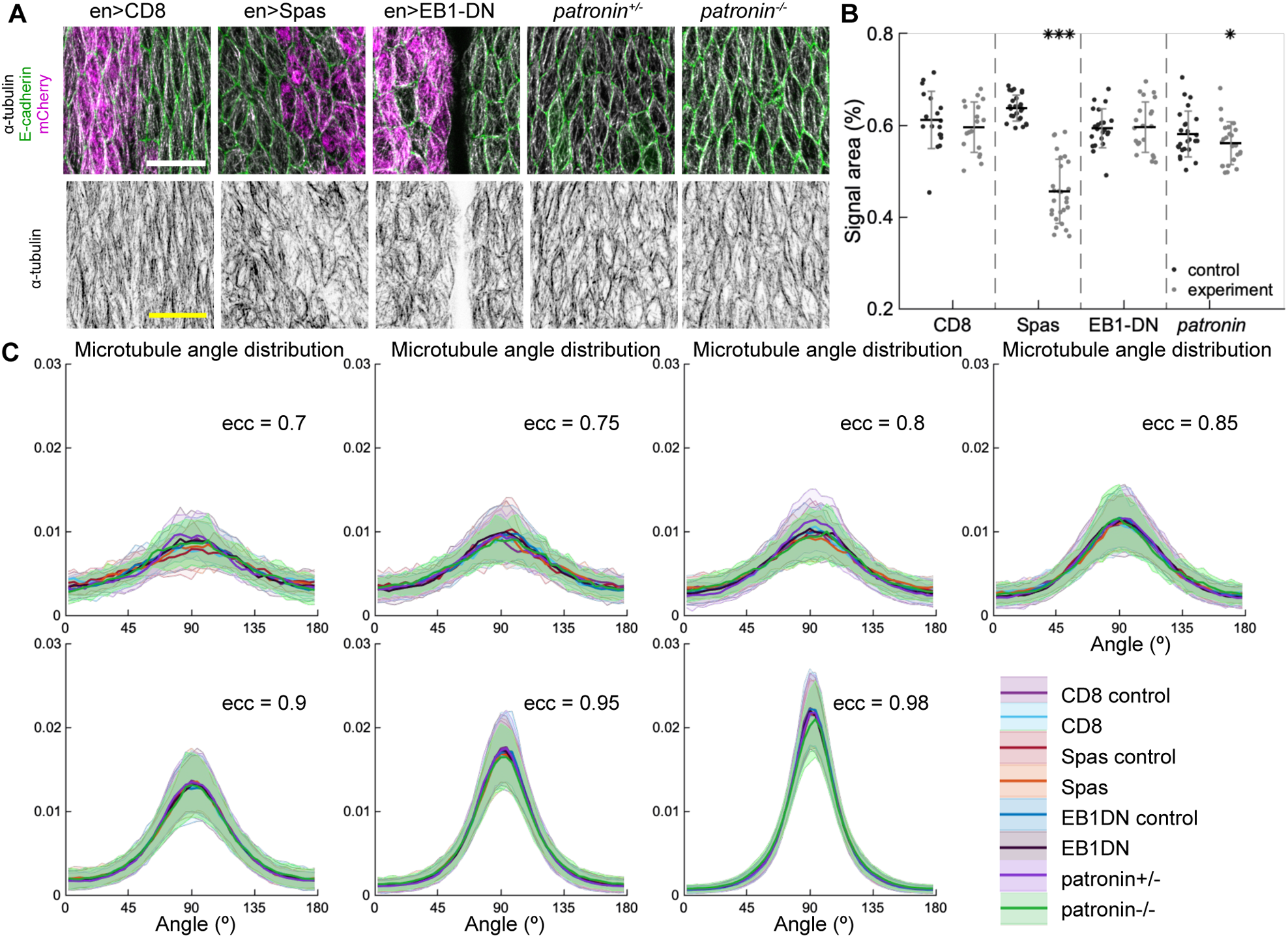
Changes to microtubule dynamics and stability do not affect their alignment. (*A*) Apical view of epidermis from conrtrol embryos and with altered microtubules. Left-to-right: embryos with CD8-Cherry (control), Spastin (Spas), and EB1-DN expressed using *paired∷*Gal4, heterozygous *Patronin*^-/+^, and homozygous *Patronin*^-/-^ embryos. Cells expressing CD8-Cherry and EB1-DN are visualized by direct fluorescence of mCherry directly fused to respective proteins, whereas cells expressing Spastin are visualized by coexpression of CD8-Cherry (*magenta*, top row). Cell outlines were visualized by immunostaining against E-cadherin (*green*, top row), and microtubules by immunostaining against *α*-Tubulin (*white*, top row; *black*, bottom row). Embryos were imaged across all developmental stages between stage 12 and 15. Scale bar - 10 *µm*. The area with no microtubules in the example of EB1-DN expression corresponds to a segmantal groove, where the cells are out of imaging focus. (B) Quantification of microtubule density in each genotype. Internal controls (cells not expressing *paired*∷Gal4) were used for CD8-Cherry, Spastin, and EB1-DN overexpression. For *Patronin*, heterozygous and homozygous embryos were compared. *** - *p* < 0.0001, * - *p* < 0.05 in comparison to respective control. (C) The microtubule angle distributions for each eccentricity (±0.025) do not significantly differ between all genotypes and relatively to controls. The distributions are shown as mean (solid line) with standard deviation (shading). blue For each eccentricity, the displayed experimental distribution is the mean distribution averaged across cells with the set eccentricity (±0.0025 for *ecc* = 0.7 − 0.95 and ±0.0005 for *ecc* = 0.98). The number of cells per eccentricity per genotype ranged from 24 to 515.

