## Supplementary Figure 1 for "Robustness of bidirectional microtubule network self-organization"

### Robustness of bidirectional microtubule network self-organization
### Supplementary information

Aleksandra Z. Płochocka,[1] Alexander M. Davie,[2] Natalia. A. Bulgakova,[3,*] and Lyubov Chumakova[2,†]

[1] *Center for Computational Biology and Center for Computational Mathematics, Flatiron Institute, New York, NY, USA, 10010*
[2] *Maxwell Institute for Mathematical Sciences, School of Mathematics, The University of Edinburgh, Edinburgh, UK, EH9 3FD*
[3] *Department of Biomedical Science, The University of Sheffield, Sheffield, UK, S10 2TN*

*
†

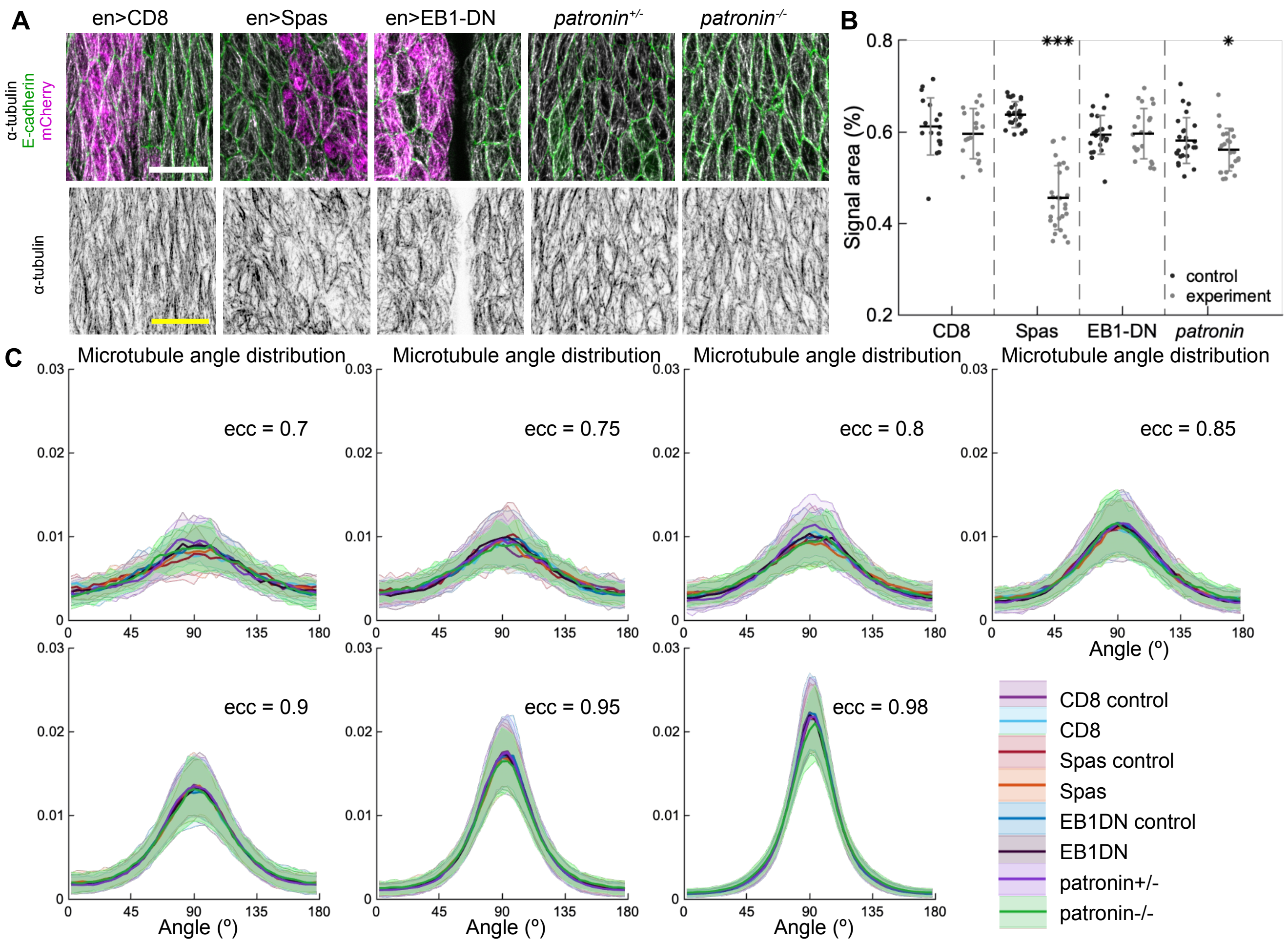


FIG. 1. Changes to microtubule dynamics and stability do not affect their alignment. (A) Apical view of epidermis from conrtrol embryos and with altered microtubules. Left-to-right: embryos with CD8-Cherry (control), Spastin (Spas), and EB1-DN expressed using *paired*::Gal4, heterozygous *Patronin*$^{-/+}$, and homozygous *Patronin*$^{-/-}$ embryos. Cells expressing CD8-Cherry and EB1-DN are visualized by direct fluorescence of mCherry directly fused to respective proteins, whereas cells expressing Spastin are visualized by coexpression of CD8-Cherry (*magenta*, top row). Cell outlines were visualized by immunostaining against E-cadherin (*green*, top row), and microtubules by immunostaining against $\alpha$-Tubulin (*white*, top row; *black*, bottom row). Embryos were imaged across all developmental stages between stage 12 and 15. Scale bar - 10 $\mu m$. The area with no microtubules in the example of EB1-DN expression corresponds to a segmantal groove, where the cells are out of imaging focus. (B) Quantification of microtubule density in each genotype. Internal controls (cells not expressing *paired*::Gal4) were used for CD8-Cherry, Spastin, and EB1-DN overexpression. For *Patronin*, heterozygous and homozygous embryos were compared. *** - $p < 0.0001$, * - $p < 0.05$ in comparison to respective control. (C) The microtubule angle distributions for each eccentricity ($\pm$ 0.025) do not significantly differ between all genotypes and relatively to controls. The distributions are shown as mean (solid line) with standard deviation (shading). blue For each eccentricity, the displayed experimental distribution is the mean distribution averaged across cells with the set eccentricity ($\pm 0.0025$ for $ecc = 0.7 - 0.95$ and $\pm 0.0005$ for $ecc = 0.98$). The number of cells per eccentricity per genotype ranged from 24 to 515.
